# Solid-state NMR Reveals Mobility-Based Organisation of the *Schizosaccharomyces pombe* Cell Wall

**DOI:** 10.64898/2026.07.20.739653

**Authors:** Ananya Singh, Teresa Massam-Wu, Mohan Balasubramanian, Wing Ying Chow

## Abstract

Fungal cell walls are hierarchically organised polysaccharide networks whose mechanical and functional properties depend on both chemical composition and molecular organisation. Although glucan synthases are essential for cell wall biosynthesis, how individual synthases shape the supramolecular architecture and dynamics of intact walls remains poorly understood. Here, we combine mobility-resolved ^13^C solid-state NMR spectroscopy with targeted genetic perturbation of the glucan synthases Ags1, Bgs1, and Bgs4 to determine how synthase activity governs the molecular organisation of the *Schizosaccharomyces pombe* cell wall. We first establish a molecular level reference for the wild-type wall by identifying the major glucan and mannan environments and resolving polysaccharides according to their mobility directly in intact cells. The rigid wall scaffold is dominated by unbranched β-1,3-glucan and α-1,3-glucan, whereas branched glucans and mannans occupy more dynamic molecular environments. Comparison with thermosensitive glucan synthase mutants reveals distinct, mutation-dependent reorganisation of both the rigid structural scaffold and the mobile polysaccharide matrix. Quantitative analysis further shows that, despite retaining broadly similar glucan compositions, the Ags1, Bgs1, and Bgs4 mutants redistribute carbohydrates among rigid, intermediate, and mobile molecular environments in distinct ways. These mutation-specific mobility fingerprints demonstrate that glucan synthases regulate not only polysaccharide biosynthesis but also how cell wall polymers are assembled, packed, and dynamically organised within the intact wall. More broadly, our findings establish molecular mobility as a sensitive signature of cell wall architecture that reveals structural consequences of biosynthetic perturbation not apparent from composition alone. Mobility-resolved solid-state NMR therefore provides a powerful framework for linking genetic perturbations to molecular dynamics and supramolecular organisation in intact fungal cell walls.

## Introduction

The fungal cell wall is a dynamic, hierarchically organised polysaccharide network that maintains cell shape, withstands internal turgor pressure, supports morphogenesis and cytokinesis, and mediates interactions with the extracellular environment.[1–4] These functions emerge not simply from the abundance of individual polysaccharides, but from their spatial organisation, intermolecular interactions, and molecular dynamics within the intact wall.[5–7] Defining how biosynthetic enzymes establish this higher order architecture is therefore central to understanding how fungal cells construct mechanically robust yet continuously remodelled extracellular assemblies.[3, 8, 9]

In the fission yeast *Schizosaccharomyces pombe* (*S. pombe*), the cell wall is composed predominantly of β-1,3-glucan, α-1,3-glucan, β-1,6-glucan, and mannans, which form structurally and functionally distinct components of the wall.[10–12] β-1,3-glucan is a major load-bearing polysaccharide and an essential component of the septum, whereas α-1,3-glucan contributes critically to lateral wall integrity, cell shape, and mechanical stability.[12–14] Branched glucans and mannans further contribute to the organisation and connectivity of the polysaccharide matrix.[15, 16] Recent studies have begun to reveal that fungal wall architecture is established through coordinated polymer synthesis and post-synthetic remodelling rather than polysaccharide production alone. In *S. pombe*, α-1,3-glucan remodelling by GH13-domain enzymes has been shown to critically influence wall architecture,[17] while branched β-1,6-glucan assembly was recently shown to involve a synthase-remodeller module in which Bgs3 functions together with the glycoside hydrolase Ghs2.[18] Collectively, these findings highlight an emerging view of the fungal cell wall as an actively constructed supramolecular assembly whose properties depend on the coordinated synthesis, modification, and organisation of its constituent polymers.

The biosynthesis of the major structural glucans in *S. pombe* is mediated by spatially and functionally specialised glucan synthases. Ags1 is required for α-1,3-glucan synthesis and contributes predominantly to lateral wall construction, whereas the β-1,3-glucan synthases Bgs1 and Bgs4 perform distinct functions during wall growth and cytokinesis.[13, 14, 19, 20] Bgs1 is particularly important for primary septum formation, while Bgs4 contributes extensively to lateral wall growth and secondary septum assembly.[13, 19] Consistent with these specialised roles, perturbation of individual synthases produces characteristic defects in cell morphology, septation, and wall integrity, demonstrating that their functions are not interchangeable.[14] However, these phenotypes do not reveal how disrupting a specific biosynthetic pathway propagates through the surrounding polysaccharide network. In particular, whether individual glucan synthases determine only the abundance and localisation of their products or also control the packing, mobility, and higher order organisation of other wall polymers remains unresolved.

Much of the current molecular picture of fungal cell walls has been established through biochemical fractionation, chemical analysis, fluorescence microscopy, immunolabelling, and electron microscopy.[1, 10, 11, 21] These approaches have provided essential information on wall composition, polymer localisation, morphology, and ultrastructure. Nevertheless, biochemical extraction disrupts native intermolecular interactions and removes polysaccharides from their cellular context, whereas imaging approaches generally lack the molecular resolution required to distinguish chemically similar polysaccharides according to their conformational environments and dynamics. Consequently, an important gap remains between identifying which polysaccharides are present and understanding how they are organised and dynamically partitioned within the intact wall.

Solid-state nuclear magnetic resonance (NMR) spectroscopy provides a complementary route to bridge this gap by enabling molecular-level investigation of complex biological assemblies without requiring polysaccharide extraction or chemical disruption.[22] In particular, ^13^C solid-state NMR has emerged as a powerful approach for investigating plant and fungal cell walls, providing information on polysaccharide composition, molecular conformation, intermolecular contacts, hydration, dynamics, and supramolecular organisation.[7, 23–27] Dynamics-based spectral editing is especially valuable for heterogeneous polysaccharide assemblies as Cross Polarisation (CP), Insensitive Nuclei Enhanced by Polarisation Transfer (INEPT), and Direct Polarisation (DP) experiments preferentially detect molecular environments with different motional properties.[28– 30] When combined with two-dimensional correlation spectroscopy, these measurements enable chemically resolved polysaccharide environments to be related directly to their molecular mobility within intact biological samples.

The ability of solid-state NMR to reveal organisational features that are inaccessible from composition alone has been demonstrated particularly clearly in fungal cell walls. Studies of *Aspergillus fumigatus* revealed a rigid, interconnected polysaccharide scaffold dominated by β-1,3-glucan and chitin, together with more dynamic matrix polysaccharides, providing a molecular framework for understanding the supramolecular architecture of filamentous fungal walls.[7, 23] More recently, solid-state NMR has begun to be integrated with defined genetic perturbations to investigate how specific biosynthetic and remodelling pathways shape cell wall architecture. In *S. pombe*, Jacob et al. demonstrated that perturbation of the α-glucan biosynthetic and remodelling machinery produces substantial changes in glucan organisation and cell wall architecture, establishing a direct connection between enzymatic function and the molecular structure of the intact wall.[17] Together, these studies establish solid-state NMR as a powerful approach for resolving molecular architecture and detecting structural reorganisation arising from defined perturbations of fungal cell wall metabolism. However, how the major, functionally specialised glucan synthases that construct different regions of the *S. pombe* cell wall differentially influence the mobility and supramolecular organisation of the surrounding polysaccharide network remains unresolved. In particular, a systematic comparison of Ags1, Bgs1, and Bgs4 perturbations within a common mobility-resolved solid-state NMR framework is needed to determine whether disruption of distinct glucan biosynthetic pathways produces characteristic reorganisations of rigid and mobile polysaccharide environments.

We recently demonstrated that fractional ^13^C labelling combined with mobility-resolved solid-state NMR provides a reproducible and cost-effective strategy for resolving rigid and mobile carbohydrate environments directly in intact *S. pombe* cells.[31] This approach provides the methodological foundation to move beyond description of the wild type (WT) wall and directly interrogate how defined genetic perturbations alter molecular organisation. The well-characterised and non-equivalent functions of Ags1, Bgs1, and Bgs4 therefore provide a powerful system for determining whether glucan synthases leave distinct molecular signatures within the architecture of the intact cell wall.

Here, we combine mobility-resolved ^13^C solid-state NMR spectroscopy with targeted perturbation of Ags1, Bgs1, and Bgs4 to determine how individual glucan synthases shape the molecular organisation of the *S. pombe* cell wall. We first establish an experimental framework for resolving polysaccharides according to molecular mobility directly in intact cells. We then use complementary two-dimensional correlation experiments to compare the rigid and mobile polysaccharide networks of WT and glucan synthase mutants, revealing mutation-specific reorganisation of glucan and mannan environments. Finally, we integrate these spectroscopic observations with complementary quantitative analyses to determine how each perturbation redistributes carbohydrates among rigid, intermediate, and mobile molecular environments. Despite broadly conserved major polysaccharide components, Ags1, Bgs1, and Bgs4 perturbations generate distinct mobility fingerprints, demonstrating that molecular dynamics reveal differences in cell wall organisation that are not apparent from polysaccharide identity alone. These findings show that perturbation of functionally specialised glucan synthases differentially reshapes the supramolecular organisation of the intact fungal cell wall and establish mobility-resolved solid-state NMR as a framework for linking defined genetic perturbations to molecular dynamics and higher order organisation in complex polysaccharide assemblies.

## Results and discussion

### Mobility-Resolved Architecture of the *S. pombe* Cell Wall

The *S. pombe* cell wall is assembled by multiple glucan synthases with spatially and functionally specialised roles. Figure 1a highlights the three glucan synthases investigated in this study: Ags1, Bgs1, and Bgs4. Ags1 synthesises α-1,3-glucan and is required for lateral wall integrity, whereas the β-1,3-glucan synthases Bgs1 and Bgs4 contribute predominantly to primary septum formation and to lateral wall and secondary septum construction, respectively.[13, 14, 19, 20] These non-equivalent biosynthetic functions provide a genetically defined system for investigating whether perturbation of individual glucan synthases produces distinct changes in the molecular organisation of the intact cell wall.

**Figure 1.**
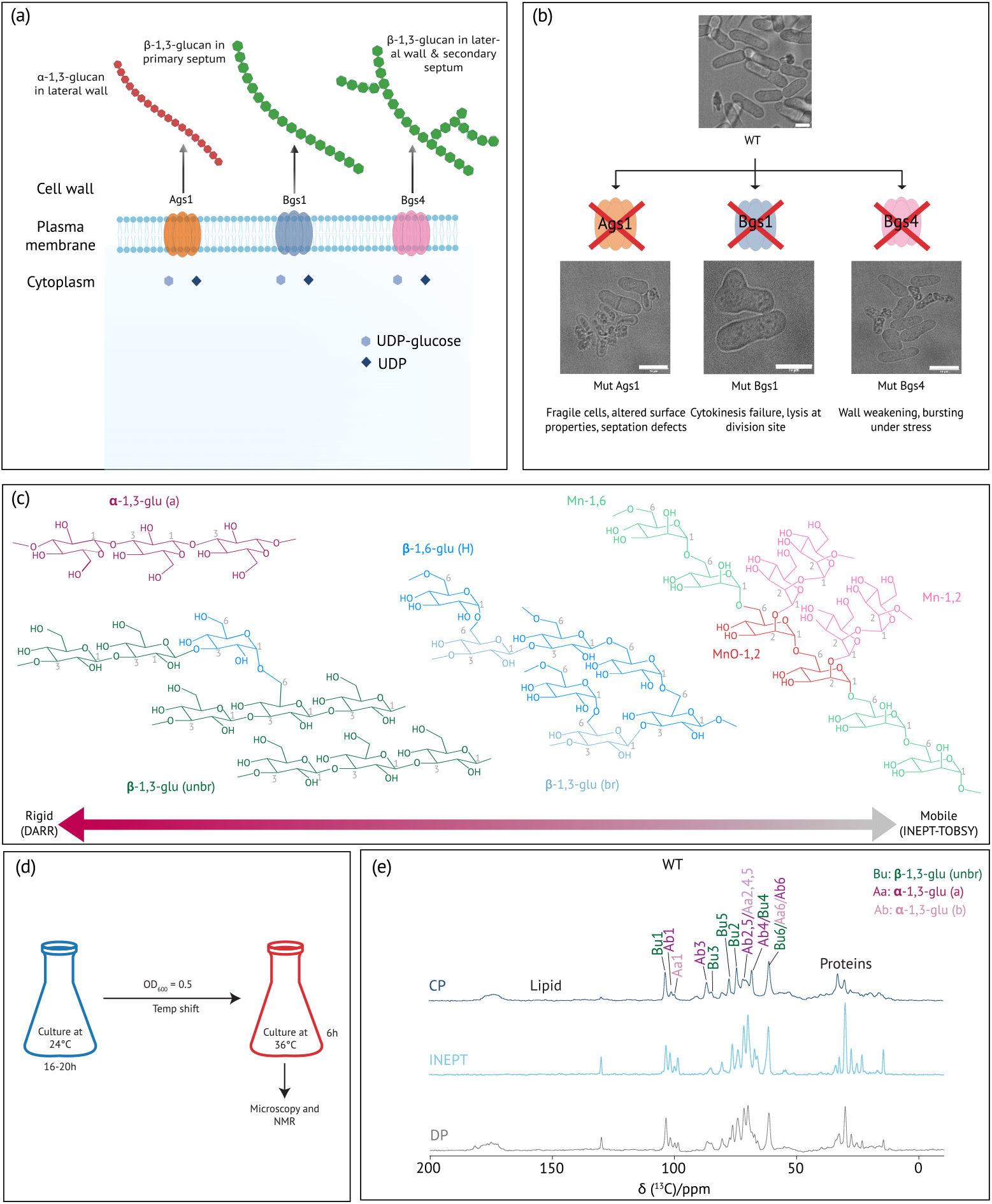
*S. pombe* has several glucan synthases, and mutations in each lead to morphological differences of the fungal cell and cell wall, which we investigated using solid-state NMR in this work. (a) Glucan synthase localisation and known functions: Ags1 synthesises α-1,3-glucan in the lateral cell wall, Bgs1 synthesises β-1,3-glucan at the primary septum, and Bgs4 synthesises β-1,3-glucan at the lateral wall and secondary septum. (b) Light microscopy of WT and mutants (Ags1, Bgs1, Bgs4) demonstrating distinct morphological defects. Scale bar: 5 *µ*m for WT and 10 *µ*m for the mutants. (c) Chemical structures of the major cell wall polysaccharides identified in this work, including: β-1,3-glucan, α-1,3-glucan, β-1,6-glucan, and galactomannan, including α-1,2-linked mannan (Mn-1,2), α-1,6-linked mannan (Mn-1,6), and O-linked α- 1,2-mannan (MnO-1,2) environments, arranged in order of molecular mobility as determined by 2D ^13^C-^13^C spectra. (d) Experimental workflow. (e) Representative ^13^C CP, INEPT, and DP spectra of WT whole cells showing rigid, mobile, and total carbohydrate contributions, respectively. All spectra were recorded at 602 MHz (^1^H) using a 3.2 mm HC Efree probe at 10 kHz MAS. Recycle delays of 2 s were used for the CP and INEPT experiments, whereas a recycle delay of 15 s was used for the DP experiment.

Consistent with their specialised roles in wall assembly, thermosensitive mutations in Ags1, Bgs1, and Bgs4 produced distinct morphological phenotypes (Figure 1b). WT cells retained the characteristic rod-shaped morphology of *S. pombe*, whereas the mutant strains exhibited pronounced alterations in cell shape and septation. These phenotypes demonstrate that disruption of individual glucan synthases compromises wall assembly in different ways. However, morphology alone cannot establish whether these defects arise from changes in polysaccharide abundance, molecular packing, dynamics, or compensatory reorganisation of the surrounding wall network.

We therefore used mobility-resolved ^13^C solid-state NMR spectroscopy to investigate the molecular organisation of the cell wall directly in intact whole cells. The major polysaccharides resolved in this study comprise β-1,3-glucan, α-1,3-glucan, branched β-1,6-glucan, and galactomannan, including α-1,2-linked mannan (Mn-1,2), α-1,6-linked mannan (Mn-1,6), and O-linked α-1,2-mannan (MnO-1,2) environments (Figure 1c). These polymers occupy distinct molecular environments within the wall and span a broad range of mobilities, from relatively rigid structural glucans to more dynamic branched glucans and mannans. Molecular mobility therefore provides an additional dimension of structural information that complements polysaccharide composition and enables the organisation of chemically heterogeneous wall components to be investigated within their native cellular environment.

The experimental workflow was designed to enable direct comparison of the glucan synthase mutants under controlled growth conditions (Figure 1d). Cells were initially cultured at 24 °C and shifted to 36 °C at an optical density (OD_600_) of approximately 0.5 to induce the thermosensitive mutant phenotypes. Following 6 h of growth at the restrictive temperature, cells were harvested for microscopy and solid-state NMR analysis. This experimental design enabled mutation-dependent morphological and molecular changes to be examined under matched growth conditions and provided intact whole cell samples for subsequent mobility-resolved NMR measurements.

Representative spectra of WT whole cells demonstrate the complementary mobility sensitivity of these experiments (Figure 1e). The CP spectrum preferentially detects rigid and motionally restricted molecular environments through strong ^1^H-^13^C dipolar couplings, whereas INEPT predominantly detects highly mobile components for which rapid molecular motion averages these interactions and permits efficient scalar-coupling-based polarisation transfer.[28, 30, 32] The DP spectrum provides broader detection of carbon environments across the cell wall and, under quantitative acquisition conditions, enables comparison of the total carbohydrate population. The substantial differences among the CP, INEPT, and DP spectral profiles demonstrate that the *S. pombe* cell wall is dynamically heterogeneous and that its polysaccharides partition into distinct mobility regimes.

Building on this mobility-resolved framework, we use two-dimensional DARR and INEPT-TOBSY spectroscopy to resolve the molecular environments of the rigid and mobile polysaccharide networks, respectively, and complementary one-dimensional analyses to quantify the redistribution of carbohydrates across mobility regimes. We first examine how perturbation of Ags1, Bgs1, and Bgs4 reorganises the rigid polysaccharide network.

### Glucan Synthase Mutations Distinctly Remodel the Rigid Cell Wall

To determine how perturbation of individual glucan synthases alters the rigid polysaccharide network, two-dimensional ^13^C-^13^C DARR spectra were acquired for WT, Ags1, Bgs1, and Bgs4 whole cell samples prepared using 10% ^13^C labelling (Figure 2). Fractional isotopic enrichment suppresses one-bond ^13^C-^13^C scalar-coupling artefacts that broaden resonances and introduce antiphase features at higher labelling densities, improving spectral resolution and line-shape quality while retaining sufficient sensitivity for comparative analysis across strains.[31, 33, 34]

**Figure 2.**
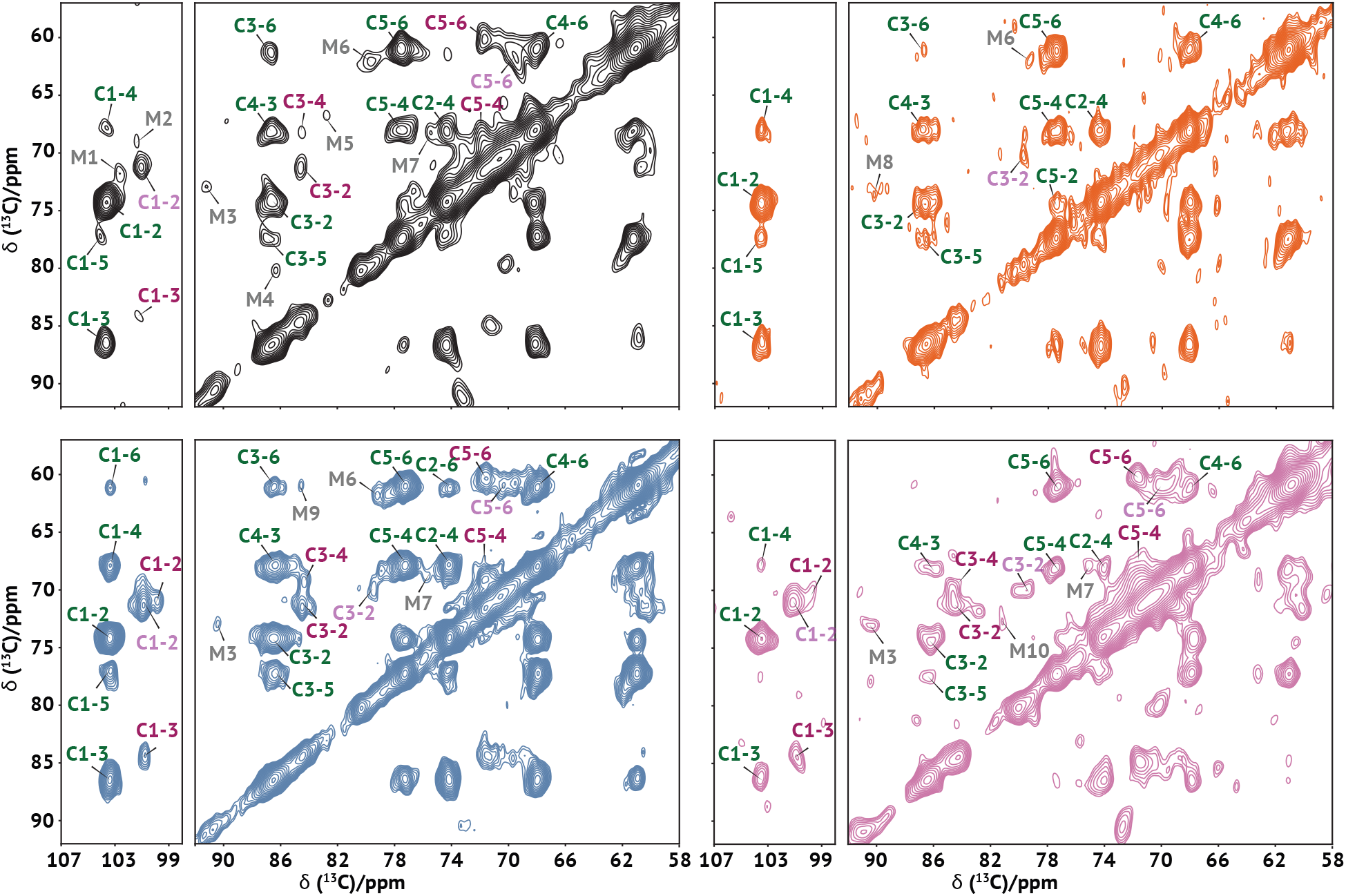
2D ^13^C-^13^C DARR spectra of WT (black), Ags1 mutant (orange), Bgs1 mutant (blue), and Bgs4 mutant (pink) *S. pombe* cell samples acquired with a 20 ms DARR mixing time at 602 MHz (^1^H) and 10 kHz MAS. The number of scans was 16 for WT, Ags1, and Bgs4 samples; 128 scans were required for the Bgs1 mutant owing to the substantially reduced rigid polysaccharide content in this strain. Additional acquisition and processing parameters are provided in the Supporting Information. The spectra selectively resolve rigid polysaccharide environments within the intact cell wall, including unbranched β-1,3-glucan (green), α-1,3- glucan (type a; dark pink), and α-1,3-glucan (type b; light pink), consistent with polymorph assignments established by Jacob et al.[17] Minor carbohydrate correlations are labelled M1-10 and summarised in Table S2.

The DARR spectra of all four strains were dominated by correlations from α-1,3-glucan and unbranched β-1,3-glucan, consistent with their preferential detection within relatively rigid molecular environments.[7, 17, 23, 31] Major correlations assigned to α-1,3-glucan environments a and b were observed together with characteristic unbranched β-1,3-glucan correlations, including C1-C2 (103.6/73.2 ppm) and C1-C3 (103.6/86.4 ppm). These dominant correlation networks were retained across WT and all three mutants, demonstrating that perturbation of Ags1, Bgs1, or Bgs4 does not eliminate the principal rigid glucan populations of the wall. Rather, the conserved assignments provide a common molecular framework within which mutation-dependent changes in relative intensity, line shape, and less populated glucan environments can be compared. In addition to the dominant glucan correlations, a series of weaker cross-peaks (M1-M10; Table S2) was resolved within the carbohydrate region. Several of these resonances, including M2 and M5, occur within chemical shift regions associated with substituted or structurally distinct glucan environments. Although the available correlations are insufficient for unambiguous assignment to specific linkage motifs, their reproducible detection across biological replicates supports their interpretation as genuine, less populated carbohydrate environments. These minor correlations therefore provide additional probes of local molecular organisation within the rigid polysaccharide network.

Comparison of the four spectra revealed distinct mutation-dependent changes superimposed on the broadly conserved glucan framework. The Ags1 mutant showed reduced α-1,3-glucan correlations together with a distinct pattern of minor carbohydrate environments: M6 (79.6/62.1 ppm) was retained from WT, M8 (90.2/73.4 ppm) was detected specifically in Ags1, whereas several WT-associated minor correlations were no longer detected under the experimental conditions. The Ags1-specific M8 correlation therefore provides additional evidence that perturbation of α-1,3-glucan biosynthesis reorganises local glucan environments within the rigid polysaccharide network rather than simply reducing the α-glucan signal. This observation is consistent with the established role of Ags1 in α-1,3-glucan biosynthesis and with recent solid-state NMR studies showing that perturbation of α-glucan biosynthesis and remodelling alters the molecular architecture of the *S. pombe* cell wall.[14, 17]

The two β-1,3-glucan synthase mutants produced distinct changes within the rigid polysaccharide network. Bgs1 retained the minor environment M3 (91.2/72.9 ppm), also detected in WT, and showed an additional substituted glucan C6 environment, M9 (84.6/60.9 ppm), that was not detected in the other strains. In contrast, Bgs4 retained M3 and the minor polysaccharide correlation M7 (75.4/68.2 ppm), but exhibited a distinct substituted glucan environment, M10 (81.2/72.7 ppm). Thus, although Bgs1 and Bgs4 both perturb β-1,3-glucan synthesis, they produce different distributions of minor glucan environments super-imposed on the broadly conserved major correlation networks. These distinct spectral fingerprints indicate that the spatially and functionally specialised roles of Bgs1 and Bgs4 result in different reorganisations of the rigid polysaccharide network.

Importantly, no major new glucan correlation networks emerged in the mutant spectra. The dominant polysaccharide assignments remained broadly conserved, whereas mutation-dependent differences were expressed primarily through changes in relative signal distributions, line shapes, and minor carbohydrate environments. Glucan synthase perturbation therefore remodels the organisation of pre-existing polysaccharide populations rather than producing wholesale replacement of the rigid wall framework.

Together, the DARR spectra reveal that Ags1, Bgs1, and Bgs4 perturbations generate distinct molecular fingerprints within a compositionally similar rigid glucan scaffold. These results indicate that individual glucan synthases influence the molecular organisation of the wall beyond determining which glucan species are present. However, dipolar-based DARR spectroscopy preferentially detects motionally restricted polysaccharides and underrepresents highly dynamic wall components because molecular motion attenuates the dipolar interactions required for efficient polarisation transfer.[22, 30] We therefore next examined whether the synthase-specific reorganisation observed in the rigid scaffold is accompanied by corresponding changes in the mobile polysaccharide network.

### Glucan Synthase Mutations Produce Distinct Mobile Wall Architectures

To determine how glucan synthase perturbation affects the dynamic polysaccharide network, two-dimensional INEPT-TOBSY spectra were acquired for WT, Ags1, Bgs1, and Bgs4 whole cell samples (Figure 3). In contrast to DARR spectroscopy, which preferentially detects motionally restricted components, INEPT-based polarisation transfer selects molecular environments with sufficient mobility to average strong dipolar interactions while retaining scalar couplings.[22, 30] The resulting spectra therefore provide a complementary view of highly mobile carbohydrate populations within intact cells.

**Figure 3.**
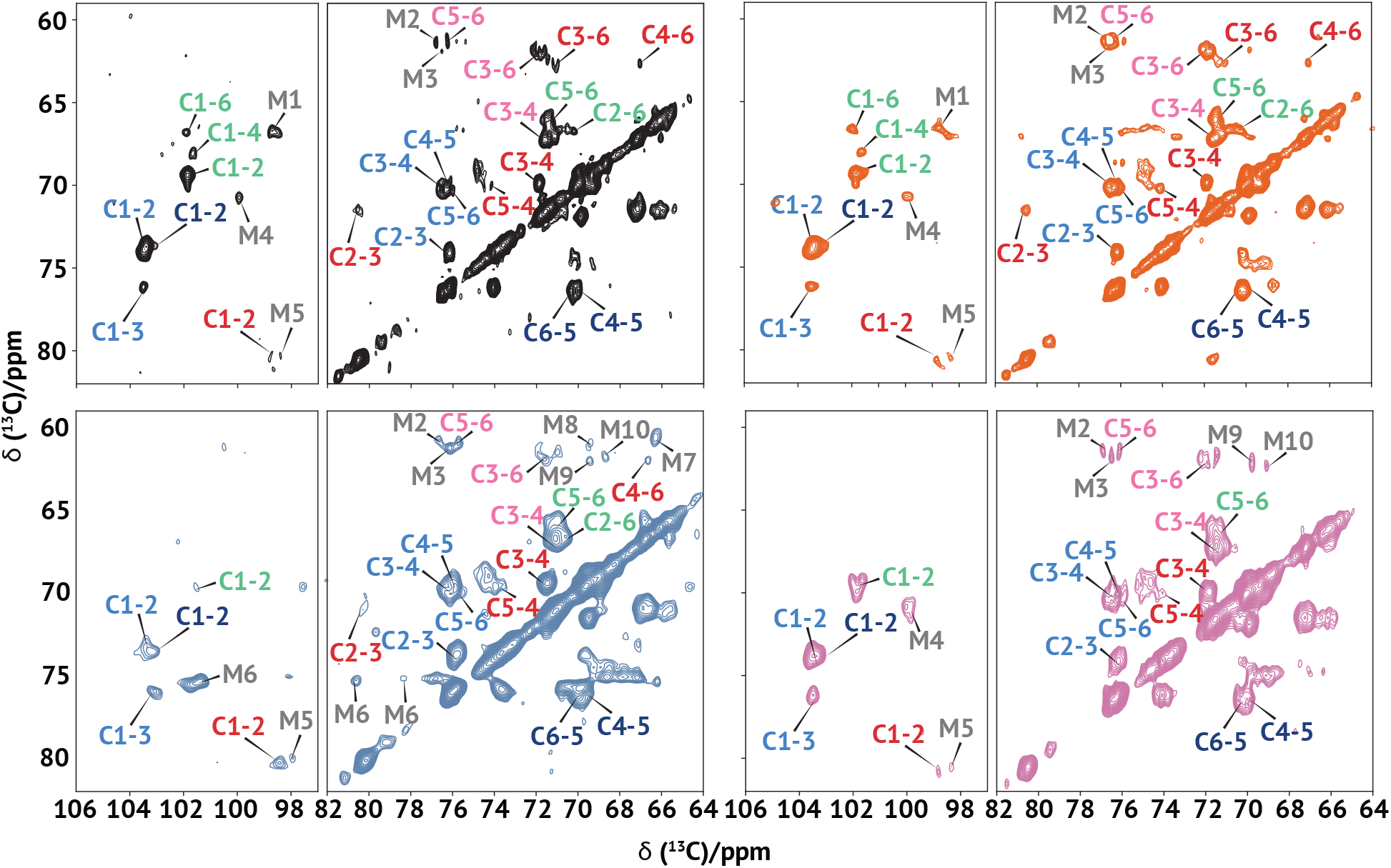
2D ^13^C-^13^C INEPT-TOBSY spectra of WT (top left, black), Ags1 mutant (top right, orange), Bgs1 mutant (bottom left, blue), and Bgs4 mutant (bottom right, pink) *S. pombe* whole cells, acquired at 10 kHz MAS with 64 scans per spectrum. WT and Ags1 spectra were recorded at ^1^H Larmor frequency of 1 GHz, whereas Bgs1 and Bgs4 spectra were acquired at 602 MHz. Additional experimental and processing parameters are provided in the Supporting Information. The spectra selectively resolve mobile polysaccharide environments within the intact cell wall. Cross-peaks are assigned to branched β-1,3-glucan (dark blue), β-1,6-glucan (light blue), α-1,2-linked mannose (pink), o-substituted α-1,2-linked mannose (red), and α- 1,6-linked mannose (green), based on characteristic fungal carbohydrate chemical shifts.[6, 7, 31] Minor carbohydrate resonances (M1-M10) are labelled and tabulated in Table S3.

The INEPT-TOBSY spectra revealed pronounced differences among the four strains. Major correlations were assigned to mobile glucan and mannan environments based on reported fungal carbohydrate chemical shifts.[7, 17, 23] In WT cells, correlations at 99.9/70.7 and 98.7/66.7 ppm were consistent with mobile mannan-associated environments, including substituted mannose residues,[6, 35] while correlations within the 69-72/61-63 ppm region were consistent with mobile carbohydrate environments involving C6 sites. Together, these signals demonstrate that the mobile fraction of the *S. pombe* cell wall contains multiple chemically distinct carbohydrate populations that are largely absent or weakly represented in the rigid DARR spectra.

Each glucan synthase mutant produced a distinct reorganisation of this mobile carbohydrate network. The Ags1 mutant exhibited an additional anomeric correlation at 104.8/71.1 ppm within the chemical shift range of β-linked carbohydrate environments.[6, 31] The appearance of this correlation in the INEPT-TOBSY spectrum indicates that perturbation of Ags1 increases the population or mobility of a β-linked glucan environment that is not resolved in WT cells. Together with the changes observed in the rigid DARR spectrum, this result demonstrates that perturbation of α-1,3-glucan biosynthesis is accompanied by reorganisation across both rigid and mobile polysaccharide populations.

The Bgs1 mutant showed the most extensive redistribution of resolved mobile carbohydrate environments, including intense correlations between 66-76 and 60-62 ppm. Peaks at 76.3/61.5 and 66.4/60.8 ppm indicate changes in highly dynamic C6-containing carbohydrate environments.[6] Several additional correlations were detected only in Bgs1, demonstrating that perturbation of this synthase generates a mobile wall fingerprint distinct from both WT and the other mutants. By contrast, Bgs4 retained several WT-like correlations but exhibited a different distribution of mobile carbohydrate signals and fewer resolved minor environments than Bgs1. Thus, perturbation of the two β-1,3-glucan synthases produces distinguishable changes in the dynamic wall network despite their shared glucan product.

The minor correlations M1-M10 (Table S3) further supported the synthase-dependent differences observed in the mobile polysaccharide network. Several environments, including M2 (76.8/61.4), M3 (76.5/61.9), and M5 (98.4/80.2), were conserved across all four strains, indicating that a subset of mobile carbohydrate populations persists despite glucan synthase perturbation. In contrast, other correlations displayed selective strain dependence: M1 (98.7/66.7)was detected only in WT and Ags1, whereas M9 69.5/62.3) and M10 (68.8/62.0) were shared by the two β-1,3-glucan synthase mutants. Bgs1 showed the most distinctive minor peak pattern, with three additional correlations (M6-M8) not detected in the other strains. Together, these differences indicate that the mutants retain common mobile carbohydrate environments while also acquiring synthase-specific spectral fingerprints, consistent with redistribution of pre-existing polysaccharides among distinct dynamic environments rather than the formation of fundamentally new carbohydrate species.

Overall, the INEPT-TOBSY spectra demonstrate that glucan synthase perturbation reorganises the dynamic polysaccharide network in a synthase-dependent manner. Importantly, the differences among Ags1, Bgs1, and Bgs4 are not restricted to the glucan populations most directly associated with each enzyme but extend across multiple mobile carbohydrate environments. Combined with the DARR spectra, these results reveal that individual synthase perturbations generate distinct molecular fingerprints across both the rigid scaffold and mobile matrix of the intact cell wall. To determine whether these spectral differences correspond to systematic redistribution of carbohydrates among mobility regimes, we next quantified the rigid, intermediate, and mobile polysaccharide populations using complementary one-dimensional NMR strategies.

### Mobility Quantification Reveals Synthase-Specific Wall Reorganisation

The DARR and INEPT-TOBSY spectra demonstrate that perturbation of Ags1, Bgs1, and Bgs4 produces distinct changes in the rigid and mobile polysaccharide networks of the *S. pombe* cell wall. However, CP and INEPT spectra alone cannot provide a common quantitative measure of wall dynamics because their signal intensities depend on different polarisation transfer mechanisms and preferentially report on motionally restricted and highly mobile molecular environments, respectively.[29, 30, 32] To compare these mobility regimes within a common analytical framework, we incorporated DP spectra as a measure of the total observable carbon population and developed two complementary quantification strategies focused on the anomeric C1 region (Figure 4a).

**Figure 4.**
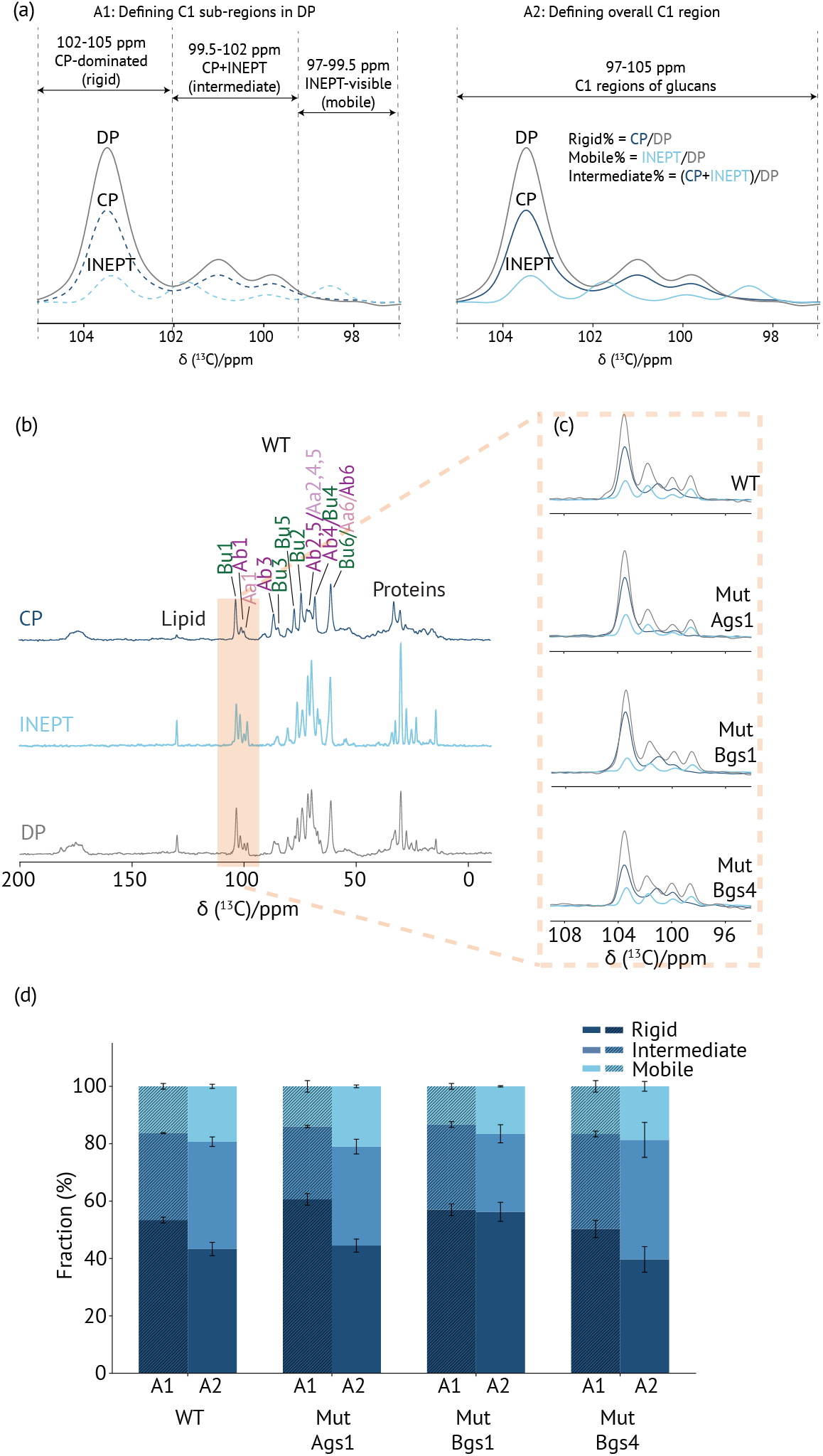
(a) NMR analysis of the anomeric C1 region (97-105 ppm) using two approaches: (A1) DP subdivision into mobility-resolved regions corresponding to rigid (CP-dominated), intermediate (visible in both CP and INEPT), and mobile (INEPT-dominated) glucans; and (A2) integration of the full C1 regions separately for CP, INEPT, and DP spectra to calculate rigid-dominated (CP/DP), mobile-dominated (INEPT/DP), and intermediate ((CP+INEPT)/DP) fractions. Spectra were scaled such that CP + INEPT intensities equal the DP intensity. (b) Representative ^13^C CP, INEPT, and DP spectra of WT whole cells showing rigid, mobile, and total carbohydrate contributions, respectively. Dominant resonances are assigned to β-1,3-glucan, α-1,3-glucan, β-1,6-glucan, and mannan components of the cell wall. All spectra were recorded at 602 MHz (^1^H) using a 3.2 mm HC Efree probe, acquisition parameters are provided in the experimental section. (c) Expanded view of the anomeric C1 region (97-105 ppm) for WT, Ags1, Bgs1, and Bgs4 strains, illustrating strain-dependent differences in peak shape and relative intensity distribution that reflect mutation-specific redistribution of carbohydrate mobility environments. (d) Quantitative comparison of rigid, intermediate, and mobile carbohydrate fractions (expressed as percentage of total C1 signal) derived from C1 spectral subdivision (A1) and CP/INEPT/DP intensity analysis (A2). Values represent mean ± SEM from three independent biological replicates (n = 3).

The glucan C1 region (105-97 ppm) was selected because it provides the greatest spectral separation of the major carbohydrate environments in the 1D ^13^C spectra while minimising overlap with the congested C4–6 region. Prior to analysis, the CP and INEPT spectra were scaled relative to the corresponding DP spectrum such that their combined signal approximately reproduced the DP spectral envelope. This empirical normalisation enabled the three experiments to be compared within a common framework while retaining the complementary sensitivity of CP and INEPT to molecular motion.[32, 36, 37] Because CP and INEPT transfer efficiencies are not direct measures of absolute molecular populations, the resulting fractions are interpreted as operationally defined mobility-associated populations rather than absolute concentrations of rigid and mobile polysaccharides.

In Approach 1 (A1), the DP C1 envelope was subdivided into three chemical shift regions according to their predominant visibility in the mobility-selective spectra (Figure 4a). The 105-102 ppm region was enriched in CP intensity and was therefore classified as rigid-dominated, the 102-99.5 ppm region contained substantial contributions from both CP and INEPT and was classified as mixed or intermediate, and the 99.5-97 ppm region was predominantly detected by INEPT and was classified as mobile-dominated. This analysis quantifies how the total DP-detected C1 signal is distributed among spectral regions associated with different motional regimes and therefore reports on redistribution of carbohydrate environments within the anomeric envelope.

In Approach 2 (A2), the entire 105-97 ppm C1 region was analysed as a single carbohydrate envelope and the relative contributions of the mobility-selective spectra were evaluated against the DP signal. CP/DP, INEPT/DP, and the remaining contribution within the normalised DP envelope were used to estimate rigid-, mobile-, and intermediate-enriched populations, respectively (Figure 4a). In contrast to the chemical shift partitioning used in A1, this analysis reports directly on differences in spectral visibility arising from molecular motion. The two strategies therefore probe complementary properties of the wall: A1 measures re-distribution among mobility-associated chemical shift regions, whereas A2 measures changes in the motional behaviour of the total glucan-associated C1 population.

Quantification across three independent biological replicates revealed distinct strain-dependent mobility profiles (Figure 4b). WT cells displayed a distributed population of carbohydrate environments across the three mobility regimes. In A1, the C1 envelope comprised (50 *±* 1)% rigid-dominated, (30 *±* 0.2)% intermediate, and (20 *±* 1)% mobile-dominated signal (Table 1). A2 similarly resolved substantial rigid- and intermediate-enriched populations, each contributing approximately 40%, together with a smaller mobile population of approximately 20% (Table 2). These distributions provide a reference mobility profile for the intact WT wall against which the glucan synthase mutants can be compared.

**Table 1.** Relative fractions of quantitative DP intensity within the three C1 chemical shift windows defined in A1. Values represent mean ± SEM from three independent biological replicates.

| **Samples** | **105-102 ppm**<br>**rigid-associated** | **102-99.5 ppm**<br>**intermediate** | **99.5-97 ppm**<br>**highly mobile** |
| --- | --- | --- | --- |
| WT | $0.5 \pm 0.01$ | $0.30 \pm 0.002$ | $0.2 \pm 0.01$ |
| Mut Ags1 | $0.6 \pm 0.02$ | $0.25 \pm 0.004$ | $0.1 \pm 0.02$ |
| Mut Bgs1 | $0.6 \pm 0.02$ | $0.3 \pm 0.01$ | $0.1 \pm 0.01$ |
| Mut Bgs4 | $0.5 \pm 0.03$ | $0.3 \pm 0.01$ | $0.2 \pm 0.02$ |

**Table 2.** Mobility-associated polysaccharide contributions estimated using A2 from integration of the C1 region in CP, INEPT, and DP spectra. CP and INEPT integrals were normalised to the corresponding DP signal to estimate the relative fractions of rigid, mobile, and intermediate-mobility polysaccharides. Values represent mean ± SEM from three independent biological replicates.

| **Samples** | **CP fraction** | **Intermediate fraction** | **INEPT fraction** |
| --- | --- | --- | --- |
| WT | $0.4 \pm 0.02$ | $0.4 \pm 0.02$ | $0.2 \pm 0.01$ |
| Mut Ags1 | $0.4 \pm 0.02$ | $0.3 \pm 0.03$ | $0.21 \pm 0.005$ |
| Mut Bgs1 | $0.6 \pm 0.03$ | $0.3 \pm 0.03$ | $0.17 \pm 0.002$ |
| Mut Bgs4 | $0.4 \pm 0.04$ | $0.2 \pm 0.02$ | $0.4 \pm 0.06$ |

Ags1 perturbation altered the wall mobility profile in both analytical frameworks, although the nature of the redistribution differed between the two approaches. A1 showed an increase in the rigid-dominated C1 region from 50% in WT to 60%, accompanied by a decrease in the mobile-dominated region from 20% to 10%. By contrast, A2 detected little change in the rigid-enriched fraction but revealed a decrease in the intermediate population and a small increase in the mobile fraction. These differences indicate that perturbation of Ags1 alters both the distribution of carbohydrate signals within the C1 envelope and their motional behaviour. Together with the loss and redistribution of α-1,3-glucan environments observed by DARR and the appearance of an additional mobile β-linked carbohydrate environment in the INEPT-TOBSY spectrum, these results demonstrate extensive reorganisation of the polysaccharide network following perturbation of α-1,3-glucan biosynthesis. Importantly, the absence of a corresponding decrease in the A2 rigid fraction indicates that changes in chemical shift distribution cannot be interpreted directly as proportional changes in overall wall rigidity.

Bgs1 produced the strongest shift towards motionally restricted carbohydrate environments. The rigid-dominated fraction increased from 50% in WT to 60% in A1, while A2 independently resolved the largest rigid-enriched population among the four strains, increasing from approximately 40% in WT to 60%. This increase was accompanied by reductions in the intermediate and mobile populations. The agreement between the two analytical approaches indicates that perturbation of Bgs1 produces a substantial redistribution towards less mobile carbohydrate environments. This quantitative response is consistent with the distinct DARR fingerprint and extensive set of Bgs1-specific mobile correlations observed by INEPT-TOBSY, demonstrating that increased representation of motionally restricted environments coexists with localised reorganisation of the mobile polysaccharide network. Thus, the Bgs1 wall is not simply globally immobilised; rather, it exhibits simultaneous redistribution across multiple mobility regimes.

Bgs4 exhibited the largest divergence between the two quantification approaches. In A1, the distribution of the C1 envelope remained similar to WT, with approximately 50%, 30%, and 20% of the signal assigned to rigid-dominated, intermediate, and mobile-dominated regions, respectively. In contrast, A2 revealed a pronounced increase in the mobile-enriched fraction from approximately 20% in WT to 40%, accompanied primarily by a decrease in the intermediate population. Thus, perturbation of Bgs4 substantially alters molecular dynamics without producing a corresponding redistribution of signal among the chemically defined C1 regions. The similarity of the A1 profiles therefore conceals a major change in molecular mobility that becomes apparent only when the same glucan-associated spectral envelope is examined using motion-sensitive experiments.

The different outcomes of A1 and A2 highlight an important feature of the analysis rather than a limitation of the quantitative framework. Chemical shift and molecular mobility report on distinct aspects of polysaccharide organisation: chemically similar glucan environments can retain comparable resonance distributions while differing substantially in motional behaviour. This distinction is particularly evident for Bgs4, where a WT-like distribution of C1 chemical shifts coexists with a markedly increased mobile population. Conversely, the Ags1 data demonstrate that redistribution within mobility-associated chemical shift regions does not necessarily correspond to an equivalent change in the globally integrated motion-sensitive fractions. The combination of the two approaches therefore distinguishes changes in spectral composition from changes in molecular dynamics.

Comparison across all four strains reveals that each glucan synthase perturbation generates a distinct quantitative mobility fingerprint. Ags1 alters the partitioning of carbohydrate environments and redistributes signal between intermediate and mobile populations, Bgs1 produces the strongest enrichment of motionally restricted carbohydrates, and Bgs4 generates the largest increase in mobile carbohydrate environments despite retaining a WT-like C1 chemical shift distribution. These quantitative differences are consistent with the mutation-specific spectral fingerprints independently observed in the rigid DARR and mobile INEPT-TOBSY spectra.

Together, the one- and two-dimensional NMR analyses establish that perturbation of Ags1, Bgs1, and Bgs4 does not simply alter the abundance of their respective glucan products. Instead, each perturbation differentially redistributes pre-existing polysaccharides among molecular environments with distinct chemical shifts and mobilities, generating characteristic architectures within the intact wall. The broadly conserved glucan assignments observed by DARR, combined with pronounced strain-dependent differences in the INEPT-TOBSY spectra and quantitative mobility profiles, demonstrate that similar polysaccharide compositions can support substantially different dynamic organisations. Molecular mobility therefore provides an organisational dimension of fungal cell wall architecture that is not captured by composition alone and reveals how functionally specialised glucan synthases differentially shape the supramolecular organisation of the intact *S. pombe* cell wall.

### Glucan Synthase Mutations Generate Distinct Cell Wall Architectures

The combined one- and two-dimensional solid-state NMR datasets support a model in which perturbation of Ags1, Bgs1, and Bgs4 influences not only glucan biosynthesis,[13, 38, 39] but also the higher order organisation and molecular dynamics of the intact fungal cell wall (Figure 5). Although the major polysaccharide classes remained broadly conserved across the four strains, DARR, INEPT-TOBSY, and quantitative mobility analyses revealed distinct redistribution of carbohydrate environments following perturbation of each synthase. The resulting models therefore represent experimentally derived differences in polysaccharide organisation and mobility rather than literal molecular structures or changes in absolute polymer abundance. WT cells exhibited a dynamically heterogeneous wall architecture spanning rigid, intermediate, and highly mobile molecular environments. DARR spectra showed that the motionally restricted network is dominated by unbranched β-1,3-glucan and α-1,3-glucan, whereas INEPT-TOBSY spectra preferentially resolved more dynamic glucan and mannan environments.[6, 17] This organisation is broadly consistent with solid-state NMR studies of fungal cell walls in which structurally constrained glucans form rigid wall networks alongside more dynamic matrix-associated polysaccharides.[6, 7] The WT model therefore provides a reference architecture in which chemically distinct polysaccharides are distributed across a continuum of molecular mobilities.

**Figure 5.**
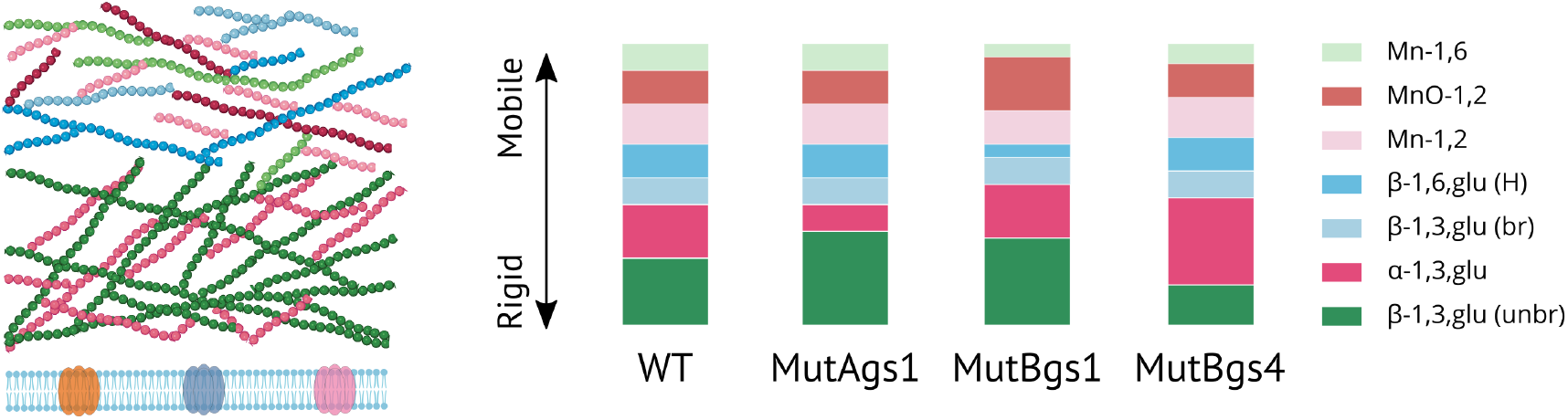
Schematic model summarising mutation-dependent reorganisation of the *S. pombe* cell wall. WT cells contain a balanced network of α-1,3-glucan, unbranched and branched β-1,3-glucan, β-1,6-glucan, and mannan-containing polysaccharides distributed across rigid and mobile environments. Ags1 perturbation reduces α-1,3-glucan and increases unbranched and branched β-1,3-glucan contributions, together with reduced β-1,6-glucan and mannan-associated environments. Bgs1 perturbation reduces unbranched and branched β-1,3-glucan and mannan-associated contributions but increases β-1,6-glucan, accompanied by the clearest redistribution towards rigid polysaccharide environments. Bgs4 perturbation reduces unbranched and branched β-1,3-glucan, β-1,6-glucan, and mannan-associated contributions, while α-1,3-glucan and O-linked mannose environments remain broadly comparable to WT. Together, the model integrates the mobility analysis based on 1D data with 2D DARR and INEPT-TOBSY spectra to show that each glucan synthase perturbation produces a distinct pattern of polysaccharide composition and molecular mobility without generating new major polysaccharide species.

Perturbation of Ags1 reorganised both the rigid and mobile polysaccharide networks. The DARR spectra revealed changes in α-1,3-glucan-associated and minor rigid carbohydrate environments, while INEPT-TOBSY detected an additional mobile β-linked carbohydrate environment. Quantitative analysis further showed redistribution among rigid-, intermediate-, and mobile-associated populations, although the two analytical approaches resolved different aspects of this response. Together, these observations indicate that perturbation of α-1,3-glucan biosynthesis propagates beyond the directly affected polymer and reorganises the broader polysaccharide network. The Ags1 model therefore depicts redistribution of pre-existing glucan environments across mobility regimes rather than simple depletion of a single structural component.

Bgs1 perturbation generated a markedly different architecture. Quantitative analysis identified the largest motionally restricted carbohydrate population among the strains examined, while the DARR spectra revealed a distinct rigid wall fingerprint and the INEPT-TOBSY spectra contained the most extensive set of strain-specific mobile carbohydrate correlations. The simultaneous increase in motionally restricted populations and emergence of distinct mobile environments demonstrates that the Bgs1 wall cannot be described simply as globally more rigid. Instead, perturbation of Bgs1 produces extensive reorganisation across multiple mobility regimes, with increased representation of rigid carbohydrate environments occurring alongside local restructuring of the dynamic polysaccharide network. The Bgs1 model therefore represents a highly redistributed wall architecture in which rigid and mobile domains are reorganised concurrently.

Bgs4 exhibited the clearest evidence that chemical shift distribution and molecular dynamics can vary independently. The DP-based subdivision of the C1 region produced a profile similar to WT, indicating broadly conserved distributions of glucan-associated chemical environments. In contrast, motion-sensitive quantification revealed the largest mobile carbohydrate population among the four strains, accompanied by a reduction in intermediate environments. These quantitative changes were consistent with the distinct INEPT-TOBSY fingerprint of the Bgs4 mutant. Thus, perturbation of Bgs4 substantially alters the physical state of the wall without producing an equivalent redistribution of the major C1 chemical shift populations. The Bgs4 model therefore illustrates how chemically similar glucan populations can adopt substantially different dynamic organisations within the intact wall.

Comparison of the four models reveals a common principle underlying the glucan synthase perturbations. None of the mutants generated fundamentally new major polysaccharide correlation networks. Instead, each perturbation redistributed pre-existing carbohydrate populations among distinct molecular environments and mobility regimes. Ags1 perturbation reorganised the α-glucan-containing network together with mobile β-linked environments, Bgs1 produced the strongest enrichment of motionally restricted populations alongside extensive reorganisation of mobile carbohydrates, and Bgs4 generated the largest increase in molecular mobility despite retaining a broadly WT-like C1 chemical shift distribution. Individual glucan synthases therefore produce distinct architectural consequences that cannot be inferred from the chemical identities of their glucan products alone.

These findings extend the emerging molecular view of fungal cell walls as dynamically organised polysaccharide assemblies whose architecture is established through coordinated biosynthesis and remodelling.[1, 17, 40] Previous solid-state NMR studies have demonstrated that fungal polysaccharides partition into molecular environments with distinct mobilities and intermolecular interactions.[6, 7] The present results further show that targeted perturbation of functionally specialised glucan synthases produces characteristic redistributions of these environments across the intact wall.

More broadly, the models establish molecular mobility as an important organisational dimension of fungal cell wall architecture. Chemically similar glucan populations can occupy different physical states depending on the biosynthetic pathway perturbed, and substantial changes in wall dynamics can occur without corresponding changes in the major polysaccharide assignments or chemical shift distributions. By integrating genetic perturbation with mobility-resolved solid-state NMR, this work provides a framework for linking individual biosynthetic enzymes to the molecular dynamics and supramolecular organisation of intact polysaccharide assemblies.

## Conclusions

Mobility-resolved ^13^C solid-state NMR combined with targeted genetic perturbation reveals how functionally specialised glucan synthases shape the molecular organisation of the intact *Schizosaccharomyces pombe* cell wall. By integrating CP, INEPT, and DP experiments with two-dimensional DARR and INEPT-TOBSY spectroscopy, we established a molecular framework that resolves the wall according to both polysaccharide identity and molecular mobility. The wild type wall comprises a rigid network dominated by unbranched β-1,3-glucan and α-1,3-glucan together with more dynamic glucan and mannan environments distributed across intermediate and highly mobile regimes.

Perturbation of Ags1, Bgs1, and Bgs4 produced distinct reorganisations of this mobility-resolved architecture. Although the major glucan correlation networks remained broadly conserved, each mutant generated a characteristic molecular fingerprint across the rigid and mobile polysaccharide populations. Ags1 perturbation reorganised the α-glucan-containing network together with mobile β-linked carbohydrate environments, Bgs1 produced the strongest enrichment of motionally restricted populations alongside extensive reorganisation of the mobile polysaccharide network, and Bgs4 generated the largest increase in molecular mobility despite retaining a broadly wild type-like distribution of C1 chemical shift environments. Thus, perturbation of individual glucan synthases does not simply alter the abundance of their respective products but differentially redistributes pre-existing polysaccharides among molecular environments with distinct dynamics and local organisation.

The complementary quantitative analyses further reveal that chemical shift distribution and molecular mobility report on distinct dimensions of cell wall architecture. This distinction is most clearly demonstrated by Bgs4, for which a broadly conserved C1 chemical shift distribution coexists with a pronounced increase in mobile carbohydrate populations. Chemically similar glucan environments can therefore adopt substantially different physical states within the intact wall, demonstrating that polysaccharide composition alone is insufficient to define cell wall architecture.

Together, these findings extend the functional consequences of glucan synthase activity beyond polysaccharide biosynthesis and establish molecular mobility as a sensitive readout of higher-order cell wall organisation. Functionally specialised synthases generate distinct architectural consequences across the surrounding polysaccharide network, revealing how perturbation of individual biosynthetic pathways propagates through an interconnected extracellular assembly. More broadly, this work establishes mobility-resolved solid-state NMR as a framework for linking defined genetic perturbations to molecular dynamics and supramolecular organisation in intact biological materials.

## Experimental

### Preparation of ^13^C-labelled samples

*Schizosaccharomyces pombe* wild type (MBY192) and thermosensitive mutant strains Ags1 (MBY10396), Bgs1 (MBY1148), and Bgs4 (MBY13423) were maintained as glycerol stocks at -80 °C. Cells were streaked onto yeast extract agar (YEA) plates and grown under permissive conditions (30 °C for WT; 24 °C for mutants) prior to liquid culture inoculation.

For solid-state NMR analysis, three independent biological batches (n = 3) of whole cell samples were prepared. Cells were cultured in 500 mL YEA medium containing 2.5 g yeast extract, 22.5 mL adenine solution (0.5%), and 15 g total glucose. For isotopic enrichment, glucose was supplied at 10% ^13^C enrichment (13.5 g unlabelled glucose and 1.5 g uniformly ^13^C-labelled glucose; Cambridge Isotope Laboratories), as described previously.[31]

Liquid cultures were grown in 2 L Erlenmeyer flasks at 24 °C with shaking at 200 rpm (Innova 42 incubator shaker) for 16-20 h until mid-log phase (OD_600_ *≈* 0.5). Thermosensitive mutants were then shifted to 36 °C for 6 h to induce the restrictive phenotype, while WT controls were processed under the same workflow for consistency. Cells were harvested by centrifugation (1500 ×*g*, 10 min, 20 °C; Eppendorf 5810R), washed three times with sterile Milli-Q water, and stored at -20 °C prior to NMR analysis.

Before rotor packing, samples were centrifuged to remove excess water and dried under a stream of nitrogen for 10-15 min to achieve partial hydration. This level of hydration has been shown to improve spectral resolution while preserving native-like polysaccharide organisation in intact fungal cell wall preparations.[31]

### Solid-state NMR experiments

Solid-state NMR experiments were performed on Bruker Avance Neo spectrometers operating at a ^1^H Larmor frequency of 602 MHz (14.15 T) at the University of Warwick. Selected INEPT-TOBSY experiments were additionally conducted at ^1^H Larmor frequency of 1 GHz (23.5 T) at the UK High-Field Solid-State NMR National Research Facility. All experiments were conducted at 268 K (to minimise sample degradation during extended acquisition) under 10 kHz magic-angle spinning (MAS) using a 3.2 mm HCN EFree probe. ^13^C chemical shifts were externally referenced to the carbonyl signal of alanine at 177.8 ppm (TMS scale).

Samples (approximately 30 mg) were packed into 3.2 mm MAS rotors by gentle loading followed by centrifugation to ensure uniform filling. Rotors were sealed immediately after packing to maintain hydration.

A combination of 1D and 2D solid-state NMR experiments was employed to probe carbohydrate domains with different motional properties. ^13^C cross-polarisation (CP) experiments were acquired with a contact time of 1 ms and recycle delay of 2 s to selectively detect rigid and immobilised components via dipolar transfer. Refocused INEPT experiments were used to selectively detect mobile components via scalar coupling transfer. Direct polarisation (DP) experiments were acquired with a recycle delay of 15 s to achieve near-quantitative signal intensities for the dominant glucan signals under the conditions used.[31] All 1D spectra were acquired with 1024 scans.

Two-dimensional ^13^C-^13^C DARR spectra were recorded with a mixing time of 20 ms to assign rigid polysaccharide resonances. Spectra were acquired with 128 scans for the Bgs1 mutant and 16 scans for WT, Ags1, and Bgs4 samples, reflecting the substantially reduced rigid polysaccharide content of the Bgs1 strain. Mobile polysaccharide environments were characterised using 2D INEPT-TOBSY experiments with a TOBSY mixing time of 5.4-6 ms and 64 scans per spectrum. Resonance assignments were validated by comparison with literature values and entries in the Complex Carbohydrate Magnetic Resonance Database (CCMRD),[41] and with assignments established for intact *S. pombe* cell walls.[17, 31]

Spectra were processed using Bruker TopSpin 4.5.0, and signals were vertically scaled such that CP + INEPT *≈* DP to enable comparison across dynamic regimes. Integration of the anomeric carbon (C1) region (97-105 ppm) was performed in TopSpin 4.5.0. Mobility-resolved and quantitative analyses were carried out as described in Figure 1d.

## Statistical analysis

All measurements were performed on three independent biological replicates (n = 3). Data are reported as mean ± standard error of the mean (SEM). Quantitative analysis of glucan distributions was performed using chemical shift-based subdivision of the C1 region (Approach 1) and CP/INEPT/DP intensity-based estimation of mobility fractions (Approach 2). Relative intensities were obtained by integrating the C1 region in TopSpin 4.5.0, following the same analysis framework as established in our previous work.[31] Variability across replicates was assessed using standard deviation and SEM calculations.

Statistical significance between conditions was evaluated using a two-sided Student’s *t* -test [42] implemented in Python (v3.13.9) with the SciPy library (v1.16.3).[43] Data visualisation, including stacked bar plots, was performed using NumPy and Matplotlib.

## Acknowledgements

We thank the EPSRC Doctoral Training Partnership for financial support through a fellowship awarded to A.S. (EP/W524645/1, Project Reference 2737036), and the University of Warwick Research Development Fund 2023/2024 (RD23105). M.K.B. and T.M.W. acknowledge support from the Wellcome Trust Senior Investigator Award (WT101885MA) and the Human Frontier Science Program (HFSP; RGP001/2023). Most NMR experiments were performed on a 602 MHz Bruker Avance NEO spectrometer funded by BBSRC (BB/T018119/1), EPSRC, and the University of Warwick, with particular thanks to assistance from Dr. Andrew Howes. Some experiments were performed at the UK High-Field Solid-State NMR National Research Facility, funded by EPSRC and BBSRC (EP/T015063/1), including the 1 GHz instrument supported through EP/R029946/1, with particular thanks to assistance from Dr. W. Trent Franks. A.S. also thank Dr. Ishutesh Jain for assistance with microscopy measurements and discussions related to the microscopy analysis.

## Supporting information

The Supporting Information documents (ZIP) are available free of charge.

The Supporting Information PDF (SI_20260720.pdf) contains:

- ^13^C spectra of all biological replicates
- Carbohydrate ^13^C chemical shift assignments
- Minor carbohydrate cross-peak assignments from 2D spectra
- NMR acquisition and processing parameters
- Additional references

Additional supporting files include:

- Raw solid-state NMR datasets
- Excel spreadsheet containing the raw quantitative data
- Python scripts for data processing, analysis, and figure generation

Some journals require a graphical entry for the Table of Contents. This should be laid out “print ready” so that the sizing of the text is correct.

The space available depends on the journal: J. Am. Chem. Soc. allows 3.25 in by 1.75 in and requires sanserif text. Some journals want different sizes: you can easily adjust here.

The two rules either side of the content are there to help judge the height of your material: they may be deleted once not required.

