## Supplementary Information for "Solid-state NMR Reveals Mobility-Based Organisation of the *Schizosaccharomyces pombe* Cell Wall"

### Availability of the raw data

The raw NMR data will be deposited on FigShare on acceptance of the manuscript.

### $^{13}$C Solid-State NMR Spectra of *S. pombe* WT and Mutant Strains

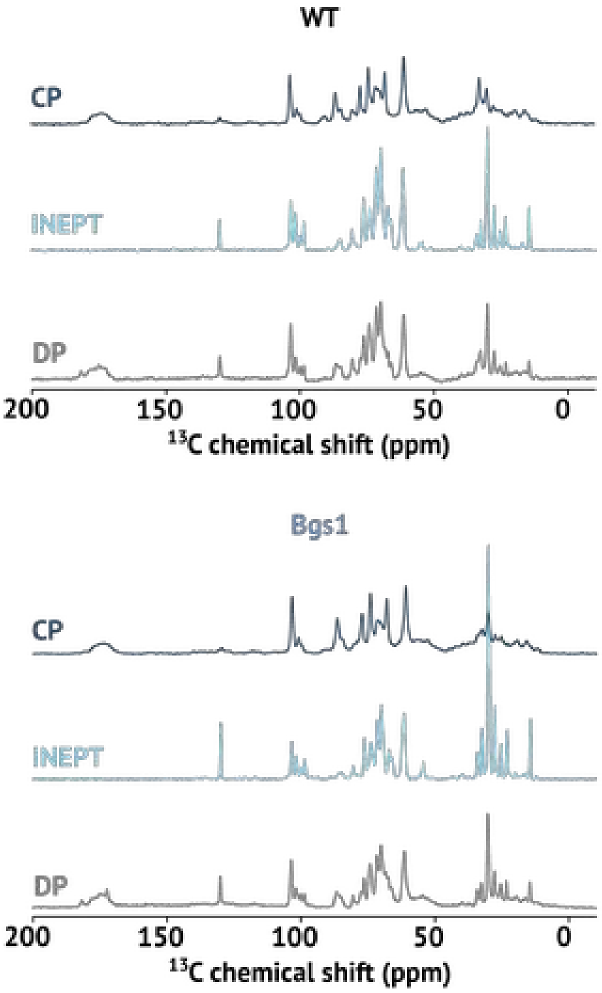

Figure S1. 1D 13C CP (dark blue), INEPT (light blue), and DP (grey) spectra of S. pombe wild type (WT) and mutant strains (Ags1, Bgs1, and Bgs4). CP spectra highlight rigid cell wall components, INEPT spectra show mobile constituents, and DP spectra provide a quantitative overview of the overall glucan content. Spectra were acquired at a 1H Larmor frequency of 602 MHz using 1024 scans and a MAS rate of 10 kHz.

### C1 Region Comparison of 13C Solid-State NMR Spectra Across Three Biological Replicates of S. pombe WT and Mutant Strains

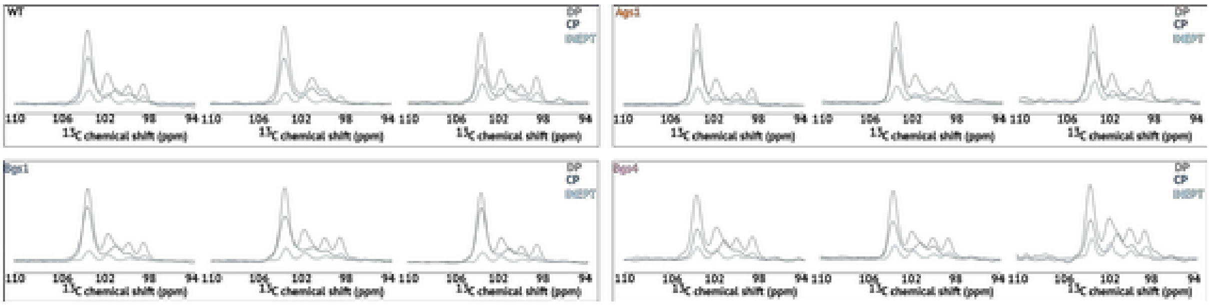

**Figure S2.** 1D $^{13}$C CP (dark blue), INEPT (light blue), and DP (grey) spectra of all the batches (n = 3) of *S. pombe* wild type (WT) and mutant strains (Ags1, Bgs1, and Bgs4), highlighting C1 regions of different glucans. Spectra are scaled such that CP and INEPT approximately add up to DP and were acquired at a $^{1}$H Larmor frequency of 602 MHz using 1024 scans and a MAS rate of 10 kHz.

### $^{13}$C chemical shifts of carbohydrates of *Schizosaccharomyces pombe*

| **Carbohydrate** | **C1** | **C2** | **C3** | **C4** | **C5** | **C6** | **Experiment** | **References** |
|---|---|---|---|---|---|---|---|---|
| α-1,3-glucan (a) | 101 | 71.4 | 84.6 | 68 | 70.1 | 60.8 | $^{13}$C-$^{13}$C DARR | Bhanja *et al.* 2014[4] |
| α-1,3-glucan (b) | 99.9 | 70.7 | 79.9 | 73.1 | 71.8 | 60.5 | $^{13}$C-$^{13}$C DARR | Jacob *et al.* 2025[5] |
| β-1,3-glucan (unbr) | 103.5 | 74.3 | 86.6 | 68.1 | 77.4 | 61.2 | $^{13}$C-$^{13}$C DARR | Shim *et al.* 2007[6] Fairweather |

| | | | | | | | | |
|---|---|---|---|---|---|---|---|---|
| | | | | | | | | *et al.* 2004[7] |
| X | 90.7 | 73.1 | 69.9 | 70 | 79.6 | 62.2 | $^{13}C$-$^{13}C$ DARR | |
| β-1,3-glucan (br) | 103 | 73.5 | | 69.5 | 76 | 70.1 | $^{13}C$-$^{13}C$ INEPT-TOBSY | Jacob *et al.* 2025[5] |
| β-1,6-glucan (H) | 103.4 | 73.8 | 76.1 | 70.1 | 74.7 | 69.0 | $^{13}C$-$^{13}C$ INEPT-TOBSY | |
| Mn-1,2 | | | 71.7 | 70.2 | 76.4 | 61.2 | $^{13}C$-$^{13}C$ INEPT-TOBSY | |
| MnO-1,2 | 98.6 | 80.4 | 71.4 | 67 | | | $^{13}C$-$^{13}C$ INEPT-TOBSY | |
| Mn-1,6 | 101.7 | 70.2 | 74.4 | 67.6 | 71.3 | 66.2 | $^{13}C$-$^{13}C$ INEPT-TOBSY | |

**Table S1.** $^{13}C$ chemical shifts of carbohydrates in *S. pombe* cell walls. "a" and "b" denote different allomorphs. Unidentified (-). Branched (Br). Reducing end (O). Unbranched (unbr). Rigid sugar resonances in $^{13}C$-$^{13}C$ DARR align with literature, while deviations in mobile sugars from $^{13}C$-$^{13}C$ INEPT-TOBSY are mainly at C4 and C5 of Mn-1,2; C4 of MnO-1,2; and C1, C2, C3, and C5 of Mn-1,6.

### Minor carbohydrate cross-peaks identified in 2D $^{13}C$-$^{13}C$ DARR spectra of WT and mutant *S. pombe* cell wall samples

| Label | Chemical shift (ppm) | Observed in | Interpretation |
|---|---|---|---|
| M1 | 102.6 / 71.8 | WT | Minor β-glucan environment |
| M2 | 101.4 /68.9 | WT | Linkage-associated β-glucan correlation |
| M3 | 91.2 / 72.9 | WT, Mut Bgs1, Mut Bgs4 | Mutant-sensitive minor glucan correlation |
| M4 | 86.3 / 80.2 | WT | Substituted/linkage-associated glucan carbon |
| M5 | 82.7 / 66.8 | WT | Possible branched glucan correlation |
| M6 | 79.6 / 62.1 | WT, Mut Ags1, Mut Bgs1 | Glucan ring-C6 correlation |
| M7 | 75.4 / 68.2 | WT, Mut Bgs1, Mut Bgs4 | Minor polysaccharide correlation |
| M8 | 90.2 / 73.4 | Mut Ags1 | Ags1-specific glucan environment |
| M9 | 84.6 / 60.9 | Mut Bgs1 | Substituted glucan C6 environment |
| M10 | 81.2 / 72.7 | Mut Bgs4 | Altered substituted glucan environment |

**Table S2.** Summary of weak and mutant-dependent carbohydrate correlations observed in the 2D $^{13}C$-$^{13}C$ DARR spectra of WT, Ags1, Bgs1, and Bgs4 cell wall samples. Peaks were labelled M1-M10 for annotation in the main manuscript figures. Tentative assignments are based on characteristic glucan chemical shift regions and comparative spectral analysis. These correlations likely reflect structurally distinct glucan linkage environments, substituted carbohydrate carbons, or intermolecular contacts within the rigid cell wall matrix.

### Minor carbohydrate cross-peaks identified in 2D $^{13}C$-$^{13}C$ INEPT-TOBSY spectra of WT and mutant *S. pombe* cell wall samples

| Label | Chemical shift (ppm) | Observed in | Interpretation |
|---|---|---|---|
| M1 | 98.7 / 66.7 | WT, Mut Ags1 | Minor mobile carbohydrate environment; likely retained following Ags1 perturbation |
| M2 | 76.8 / 61.4 | WT, Mut Ags1, Mut Bgs1, Mut Bgs4 | Conserved mobile carbohydrate environment present across all strains |
| M3 | 76.5 / 61.9 | WT, Mut Ags1, Mut Bgs1, Mut Bgs4 | Conserved mobile environment with enhanced intensity in mutants, suggesting increased mobility or exposure |
| M4 | 99.9 / 70.7 | WT, Mut Ags1, Mut Bgs4 | Dynamic carbohydrate environment enriched in Ags1 and Bgs4, indicative of wall remodelling |
| M5 | 98.4 / 80.2 | WT, Mut Ags1, Mut Bgs1, Mut Bgs4 | Conserved substituted or structurally distinct carbohydrate environment |
| M6 | 101.6 / 75.6 | Mut Bgs1 | Bgs1-specific mobile carbohydrate environment associated with septation-related wall reorganisation |
| M7 | 66.4 / 60.8 | Mut Bgs1 | Bgs1-specific mobile carbohydrate environment, possibly linked to compensatory remodelling |
| M8 | 69.5 / 61.2 | Mut Bgs1 | Minor mutant-specific carbohydrate environment indicative of altered wall heterogeneity |
| M9 | 69.5 / 62.3 | Mut Bgs1, Mut Bgs4 | Mobile carbohydrate environment associated with β-glucan synthase perturbation |
| M10 | 68.8 / 62.0 | Mut Bgs1, Mut Bgs4 | Minor dynamic carbohydrate environment enriched in β-glucan synthase mutants |

**Table S3.** Summary of weak and mutant-dependent carbohydrate correlations observed in the 2D $^{13}C$-$^{13}C$ INEPT-TOBSY spectra of WT, Ags1, Bgs1, and Bgs4 cell wall samples. Peaks were labelled M1-M10 for annotation in the main manuscript figures. Tentative interpretations are based on chemical shift position and comparative spectral analysis. These correlations likely arise from structurally distinct mobile carbohydrate environments and highlight mutation-dependent reorganisation of the dynamic cell wall network

### Solid-State NMR Acquisition and Processing Parameters

| Experiment | Sample | Spectrometer frequency | Acquisition parameters | | | | | | | | Processing parameters | |
|---|---|---|---|---|---|---|---|---|---|---|---|---|
| | | | t (h) | d1 (s) | NS | td2 | td1 | aq2 (ms) | aq1 (ms) | sw2 (ppm) | sw1 (ppm) | Wdw | Parameter |
| 1D $^{13}C$ CP | WT | 602 MHz | 35m | 2 | 1024 | | 1500 | | 20.5 | | 330.25 | Qsine | SSB 2 |
| | Mut Ags1 | 602 MHz | 35m | 2 | 1024 | | 1500 | | 20.5 | | 330.25 | Qsine | SSB 2 |
| | Mut Bgs1 | 602 MHz | 35m | 2 | 1024 | | 1500 | | 20.5 | | 330.25 | Qsine | SSB 2 |
| | Mut Bgs4 | 602 MHz | 35m | 2 | 1024 | | 1500 | | 20.5 | | 330.25 | Qsine | SSB 2 |
| 1D $^{13}C$ INEPT | WT | 602 MHz | 35m | 2 | 1024 | | 1500 | | 20.5 | | 330.25 | Qsine | SSB 2 |
| | Mut Ags1 | 602 MHz | 35m | 2 | 1024 | | 1500 | | 20.5 | | 330.25 | Qsine | SSB 2 |
| | Mut Bgs1 | 602 MHz | 35m | 2 | 1024 | | 1500 | | 20.5 | | 330.25 | Qsine | SSB 2 |
| | Mut Bgs4 | 602 MHz | 35m | 2 | 1024 | | 1500 | | 20.5 | | 330.25 | Qsine | SSB 2 |
| 1D $^{13}C$ DP | WT | 602 MHz | 4h 17m | 15 | 1024 | | 1500 | | 20.5 | | 330.25 | Qsine | SSB 2 |
| | Mut Ags1 | 602 MHz | 4h 17m | 15 | 1024 | | 1500 | | 20.5 | | 330.25 | Qsine | SSB 2 |
| | Mut Bgs1 | 602 MHz | 4h 17m | 15 | 1024 | | 1500 | | 20.5 | | 330.25 | Qsine | SSB 2 |
| | Mut Bgs4 | 602 MHz | 4h 17m | 15 | 1024 | | 1500 | | 20.5 | | 330.25 | Qsine | SSB 2 |
| 2D $^{13}C$-$^{13}C$ DARR | WT | 602 MHz | 2h 20m | 20 | 16 | 1024 | 256 | 35.0 | 4.3 | 331.68 | 199.01 | Qsine | SSB 2 |
| | Mut Ags1 | 602 MHz | 2h 20m | 20 | 16 | 2048 | 256 | 35.0 | 4.3 | 331.68 | 199.01 | Qsine | SSB 2 |
| | Mut Bgs1 | 602 MHz | 23h 21m | 20 | 128 | 2000 | 256 | 26.0 | 4.0 | 254.05 | 264.21 | GM | SSB 2 |
| | Mut Bgs4 | 602 MHz | 2h 20m | 20 | 16 | 1024 | 256 | 35.0 | 4.3 | 331.68 | 199.01 | Qsine | SSB 2 |
| 2D $^{13}C$-$^{13}C$ INEPT-TOBSY | WT | 1 GHz | 10h 57m | 2 | 64 | 4864 | 300 | 39.0 | 7.5 | 248.44 | 79.5 | GM | SSB 2 |
| | Mut Ags1 | 1 GHz | 16h 26m | 2 | 96 | 3072 | 300 | 39.0 | 7.5 | 248.44 | 79.5 | GM | SSB 2 |
| | Mut Bgs1 | 602 MHz | 19h 42 m | 2 | 64 | 2048 | 540 | 35.0 | 9.0 | 262.11 | 198.15 | GM | SSB 2 |
| | Mut Bgs4 | 602 MHz | 19h 44m | 2 | 64 | 2776 | 540 | 35.0 | 9.0 | 262.11 | 198.15 | GM | SSB 2 |

**Table S4.** Acquisition and processing parameters used for the 1D $^{13}C$ CP, INEPT, and DP experiments, and the 2D $^{13}C$-$^{13}C$ DARR and $^{13}C$-$^{13}C$ INEPT-TOBSY experiments performed on *S. pombe* wild type (WT) and mutant strains (Ags1, Bgs1, and Bgs4). Abbreviations: NS, number of scans; td1 and td2, number of points in the indirect and direct dimensions; aq1 and aq2, acquisition times in the indirect and direct dimensions; sw1 and sw2, spectral widths in the indirect and direct dimensions; WDW, window function applied during processing; SSB, sine bell shift; GM, Gaussian multiplication.
